# eVIP2: Expression-based variant impact phenotyping to predict the function of gene variants

**DOI:** 10.1101/872028

**Authors:** Alexis M. Thornton, Lishan Fang, Casey O’Brien, Alice H. Berger, Marios Giannakis, Angela N. Brooks

## Abstract

While advancements in genome sequencing have identified millions of somatic mutations in cancer, their functional impact is poorly understood. We previously developed the expression-based variant impact phenotyping (eVIP) method to use gene expression data to characterize the function of gene variants. The eVIP method uses a decision tree-based algorithm to predict the functional impact of somatic variants by comparing gene expression signatures induced by introduction of wild-type versus mutant cDNAs in cell lines. The method distinguishes between variants that are gain-of-function, loss-of-function, change-of-function, or neutral. We present eVIP2, software that allows for pathway analysis (eVIP Pathways) and usage with RNA-seq data. To demonstrate the eVIP2 software and approach, we characterized two recurrent frameshift variants in *RNF43*, a negative regulator of Wnt signaling, frequently mutated in colorectal, gastric and endometrial cancer. *RNF43 WT*, *RNF43 R117fs*, *RNF43 G659fs*, or *GFP* control cDNA were overexpressed in HEK293T cells. Analysis with eVIP2 predicted that the frameshift at position 117 was a loss-of-function mutation, as expected. The second frameshift at position 659, was, surprisingly, predicted to be a gain-of-function mutation. Additional eVIP Pathways analysis of *RNF43 G659fs* predicted 10 pathways to be significantly altered, including TNF alpha via NFKB signaling, KRAS signaling, and hypoxia. To validate these predictions, we performed reporter assays and found that all eVIP2 impactful pathways tested in the assay were activated by expression of *RNF43 G659fs*, but not by expression of RNF43 WT, supporting that *RNF43 G659fs* is a gain-of-function mutation and its effect on the identified pathways. The eVIP2 method is an important step towards overcoming the current challenge of variant interpretation in the implementation of precision medicine. eVIP2 is available at https://github.com/BrooksLabUCSC/eVIP2.

## Introduction

While advancements in genome sequencing have identified millions of somatic mutations in cancer [1–3], interpretation of these variants remains a major challenge to the implementation of precision medicine. Distinct assays are generally used to determine mutation impact for each individual gene being studied, slowing down the process of variant interpretation. Previously, no single assay could rapidly profile the functional impact of a diverse set of genes. Earlier studies demonstrated the feasibility of using gene expression signatures as “fingerprints” of molecular function [4].

In Berger et al., we presented the expression-based variant-impact phenotyping (eVIP) method that uses gene expression changes to distinguish impactful from neutral somatic mutations [5]. This study used the L1000 assay, which measures the abundance of 978 “landmark” genes [6]. eVIP was used to characterize 194 somatic mutations in 53 genes identified in primary lung adenocarcinomas, demonstrating the value of systematic functional interpretation of variants using gene expression data.

Here, we present advancements in the eVIP approach and software, called eVIP2. eVIP2 includes a more automated pipeline and improvements in user-friendliness. Also, we use the eVIP2 method with RNA-seq data, which provides a better view of the full transcriptome than L1000 data and is more widely used. In addition to functional predictions on the overall effect of each variant, we also include an approach called eVIP Pathways to predict the functional impact on specific cellular signaling pathways. We recommend using eVIP Pathways with RNA-Seq data.

We show that eVIP2 can be used to characterize variants of unknown function and identify their downstream effect on cellular signaling pathways.

## Design and Implementation

### Algorithm for expression-based variant impact phenotyping

The eVIP approach assumes that the underlying expression data were derived from cell line experiments from overexpression of WT cDNA, mutant cDNA(s), and control cDNA(s) (Fig 1A). The eVIP method uses a decision tree-based algorithm that determines the functional impact of a mutation by comparing gene expression signatures induced by wild-type and mutant ORFs (Fig 1B). The two features of gene expression changes that eVIP uses are signal strength and signature identity. The signal strength is a quantitative measure of replicate consistency (WT vs. WT or variant vs. variant). The signature identity is found by calculating the correlation between the transcripts in the wild-type signature and those in the mutant signature (WT vs. variant).

**Fig 1.**
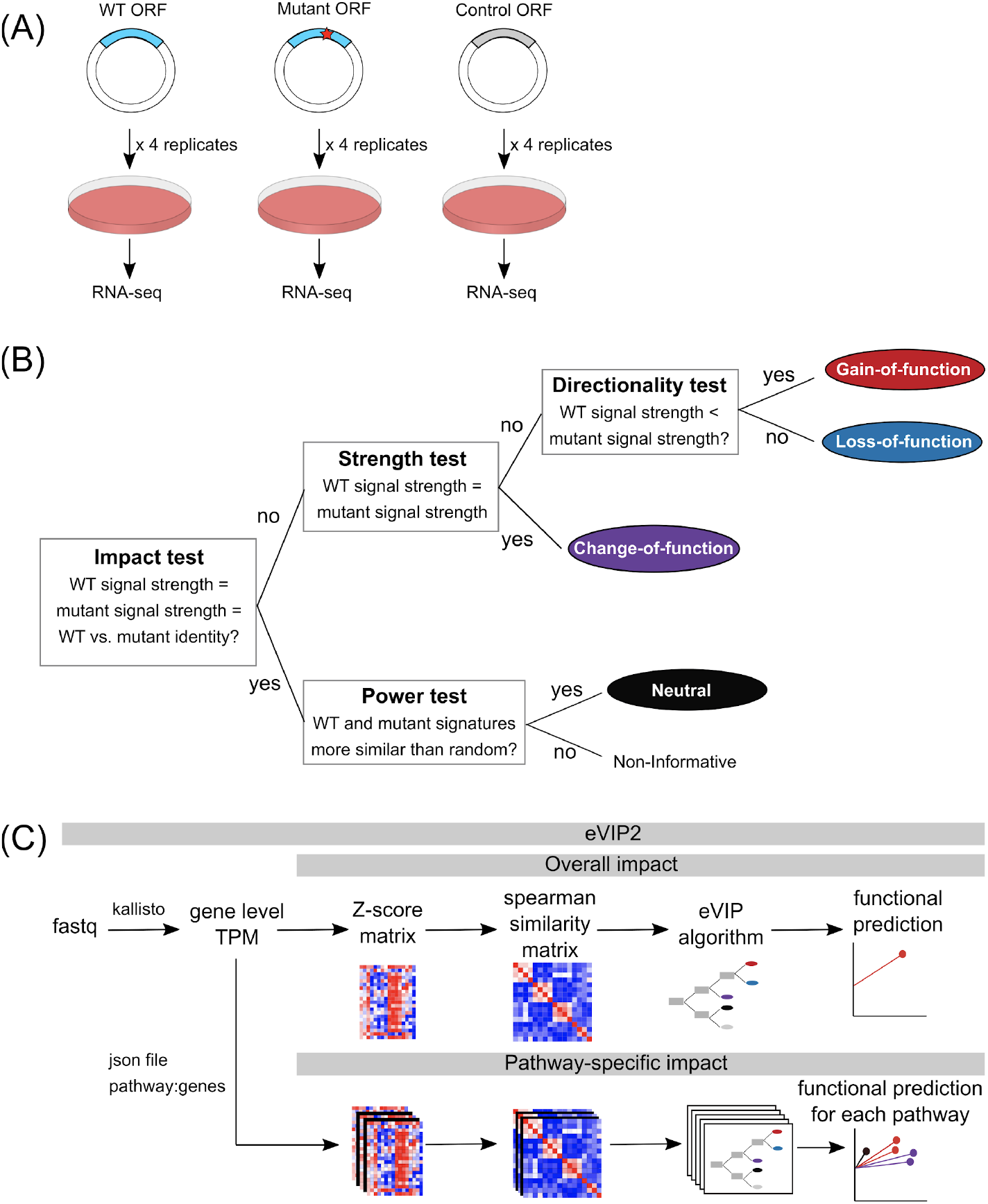
The eVIP algorithm uses RNA-seq data to predict the function of somatic mutations and pathways. (A) Overview of experimental approach (B) Schematic of the eVIP decision tree-based eVIP algorithm. The impact test is a Kruskal-Wallis test of the three distributions: wild-type replicate self-correlation, mutant replicate self-correlation, and wild-type versus mutant correlation. It outputs a Bonferonni-adjusted p-value, which represents the likelihood of mutation impact. Impactful mutations are then tested for their directional impact. For non-impactful mutations, a “power test” determines whether the two signatures are similar to one another due to a real signal or due to noise.(C) Overview of the eVIP2 pipeline which incorporates overall impact and pathway impact. The eVIP2 pipeline uses gene level counts (TPM) to predict the functional impact of a mutation. Many mutations can be processed in parallel. There is the option for eVIP Pathways, which predicts the impact of each mutation on a pathway.

At the root of the decision tree, we test the null hypothesis that the mutant signature and the wild-type signature are indistinguishable, and give the overall impact prediction p-value for the given variant. This is based on a Kruskal-Wallis test performed on three distributions: wild-type replicate self-correlation, mutant replicate self-correlation, and wild-type versus mutant correlation.

When the overall impact p-value is less than 0.05 (with multiple-testing correction), the variant is impactful. The variant can then be further characterized as a change-of-function (COF), gain-of-function (GOF), or loss-of-function (LOF). A Wilcoxon test is done to determine if there is no difference in self-replicate correlation between the mutant and the wild type. If the difference is not significant the variant is a change-of-function. If the difference is significant, we then look to find the direction of the difference in self-replicate correlation. When the mutant self-replicate correlation is less than wild-type self-replicate correlation, it has a negative impact score and it is loss-of-function. When it is greater, it has a positive impact score and the variant is a gain-of-function.

When the overall impact p-value is greater than 0.05, the variant may have no impact on gene function or be non-informative. A wilcoxon test is then done by comparing the wild-type versus mutant replicate correlation distribution to a null distribution. The null distribution is determined by comparing the mutant signature to control signature(s) and iterating this comparison 1,000 times. Thus, a variant is predicted to be neutral if the mutant signature is indistinguishable from wild-type and if the mutant signature is more similar to wild-type than control signature comparisons.

### Data processing and eVIP2 pipeline

For overall functional predictions, the eVIP2 software takes in L1000 Z-scores or RNA-seq gene level transcript per million (TPM) counts (Fig 1C). It is recommended to filter out genes with low expression and do a log2 transformation. For pathway analysis with eVIP2, raw Kallisto [7] outputs are required.

Using the log2 TPM counts, eVIP2 calculates Z-scores across the replicates in all conditions and then reduces the data to a sample by sample self-correlation matrix. It is recommended to have at least four replicates for each condition. In the original implementation using L1000 data, a weighted connectivity score was used as a measure of correlation. Using the large L1000 dataset that tested the impact of 194 somatic mutations [5], we compared the weighted connectivity score to spearman rank as a measure of correlation. We found that the corrected p-values from the impact test using either correlation method were comparable (S1 Fig); therefore, we used spearman rank correlation for RNA-Seq data. Using the correlation matrix, the described eVIP algorithm is run to give a prediction of LOF, GOF, COF, Neutral, or non-informative.

In addition to a variant receiving an overall functional impact prediction, with eVIP Pathways, the user can also determine pathway-specific functional calls, which allows for more specific functional analysis. eVIP Pathways requires a JSON file representing the mapping of pathways to their genes. Custom gene sets or curated gene sets from MsigDB, Kegg, and Reactome can be used [8–10]. A script called “create_pathway_JSON.py” is included to convert gene sets in GMT file format into JSON files that can be used with eVIP Pathways.

For pathway analysis, eVIP Pathways first finds differentially expressed genes that are specific to the WT or mutant. The WT gene and each mutant are compared to the control using DESeq2 [11]. We define mutation-specific genes as genes that are differentially expressed only in the control versus mutation and not in the control versus WT. These genes represent a new function caused by the mutant. The WT-specific genes are differentially expressed only in the control versus WT and not in the control versus mutant. These are genes that are expected to be affected by normal WT function, but are not affected by the mutant, and therefore contribute to mutant loss of function. eVIP Pathways is then run separately using the WT-specific and mutant-specific genes (with multiple-testing correction).

The standard approach to understand the function of a mutation is to look at differentially expressed genes in cells with and without induced expression of the mutant. We believe that analysis of mutant-specific differentially expressed genes allows a better discernment of mutation function, by disregarding the preserved effects of the WT gene.

## Results

### eVIP2 identifies different impacts of two frameshift mutations in *RNF43*

To demonstrate the utility of the eVIP2 approach, we examined the impact on *RNF43* gene function of its most common mutations, R117fs and G659fs (Fig 2A). *RNF43* encodes for a cell surface transmembrane E3 ubiquitin-protein ligase that acts as a negative regulator of the Wnt signaling pathway [12–14]. Recurrent *RNF43* mutations are predicted to create neopeptides [15,16]. Over 18% of colorectal adenocarcinomas and endometrial carcinomas have *RNF43* mutations [17] with p.G659Vfs*41 and p.R117Afs*41 being the most common. *RNF43* mutations are associated with microsatellite-instable tumors and are mutually exclusive with inactivating APC mutations [17].

**Fig 2.**
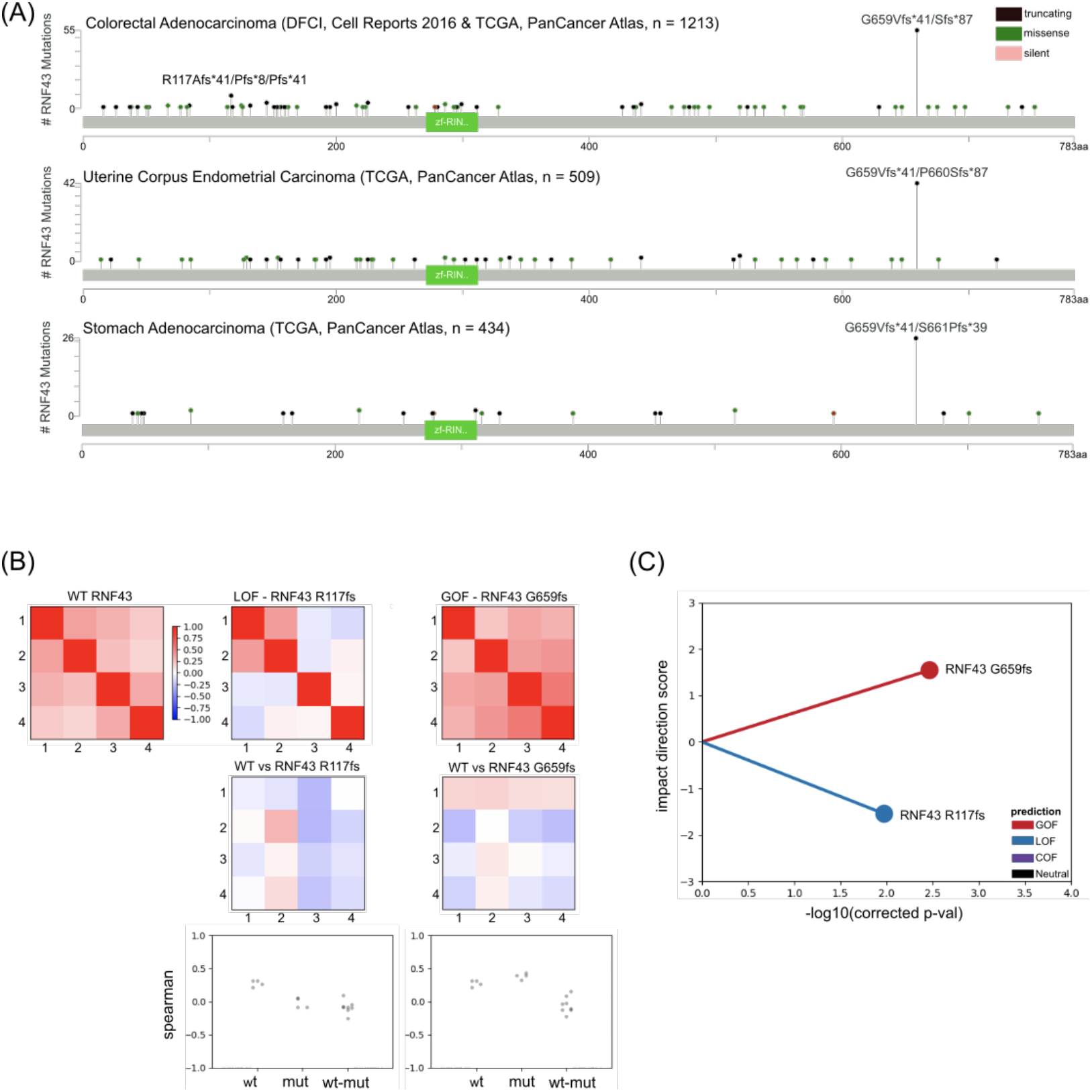
eVIP identifies functional differences between two RNF43 frameshift mutations. (A) Cbioportal “MutationMapper”[28], which shows the frequency of mutations in RNF43 in three different cohorts. The ring finger domain is indicated in green. (B) Heat map representation of WT replicate consistency (WT vs WT) or variant replicate consistency (RNF43 G659fs vs RNF43 G659fs and RNF43 R117fs vs RNF43 R117fs). Signature identity (WT vs variant) is represented by heatmaps in the second row. Dot-plot representation of replicate consistency and signature identity measured by spearman rank correlation. (C) A “sparkler” plot representation of eVIP predictions. A point represents a variant. The x axis represents the Kruskal wallis “impact test” -log10(adjusted p-value). The y axis is the “impact direction score”, the absolute value of which is equal to the –log10 (adjusted p-value) of a Wilcoxon test directly comparing wild-type and mutant ORF replicate consistency.The sign of the impact direction score is positive if the mutant replicate consistency is greater than WT and negative if the mutant replicate consistency is less than the WT replicate consistency.

Wild-type *RNF43* and both frameshift mutations were overexpressed in HEK293T cells, with 4 replicates in each condition, and expression profiling was performed using RNA-Seq. The overexpression of *RNF43 WT*, *RNF43 R117fs*, and *RNF43 G659fs*, was confirmed by inspecting the RNA-Seq reads and through Western blot validation (S2 Fig). The eVIP2 overall impact predicted the *RNF43 R117fs* variant to cause a loss of function, which is consistent with the R117fs mutation leading to a premature stop codon early in the gene, thereby disrupting the majority of the protein (Figs 2 B-C). Interestingly, the G659fs variant was predicted to cause a gain of function (Figs 2 B-C). The G659fs frameshift occurs 126 amino acids away from the end of the wild-type gene and the termination codon in the new reading frame is 41 amino acids away from the frameshift. Frameshifts that occur late in the gene can often be assumed to be loss-of-function or not alter wild-type protein function; however the hotspot mutational pattern of G659fs (Fig 2A) and our eVIP2 overall impact prediction (Figs 2 B-C,S1 File) point to a gain-of-function.

### Mutation-specific and WT-specific differentially expressed genes recapitulate overall LOF and GOF calls

We took advantage of the RNA-seq data and used eVIP Pathway analysis to find which pathways are impactful in each variant. To further examine the *RNF43* frameshift mutations, we defined “mutation-specific” and “WT-specific” gene sets (Fig 3A). This allows us to find gene expression changes specific to each frameshift.

**Fig 3.**
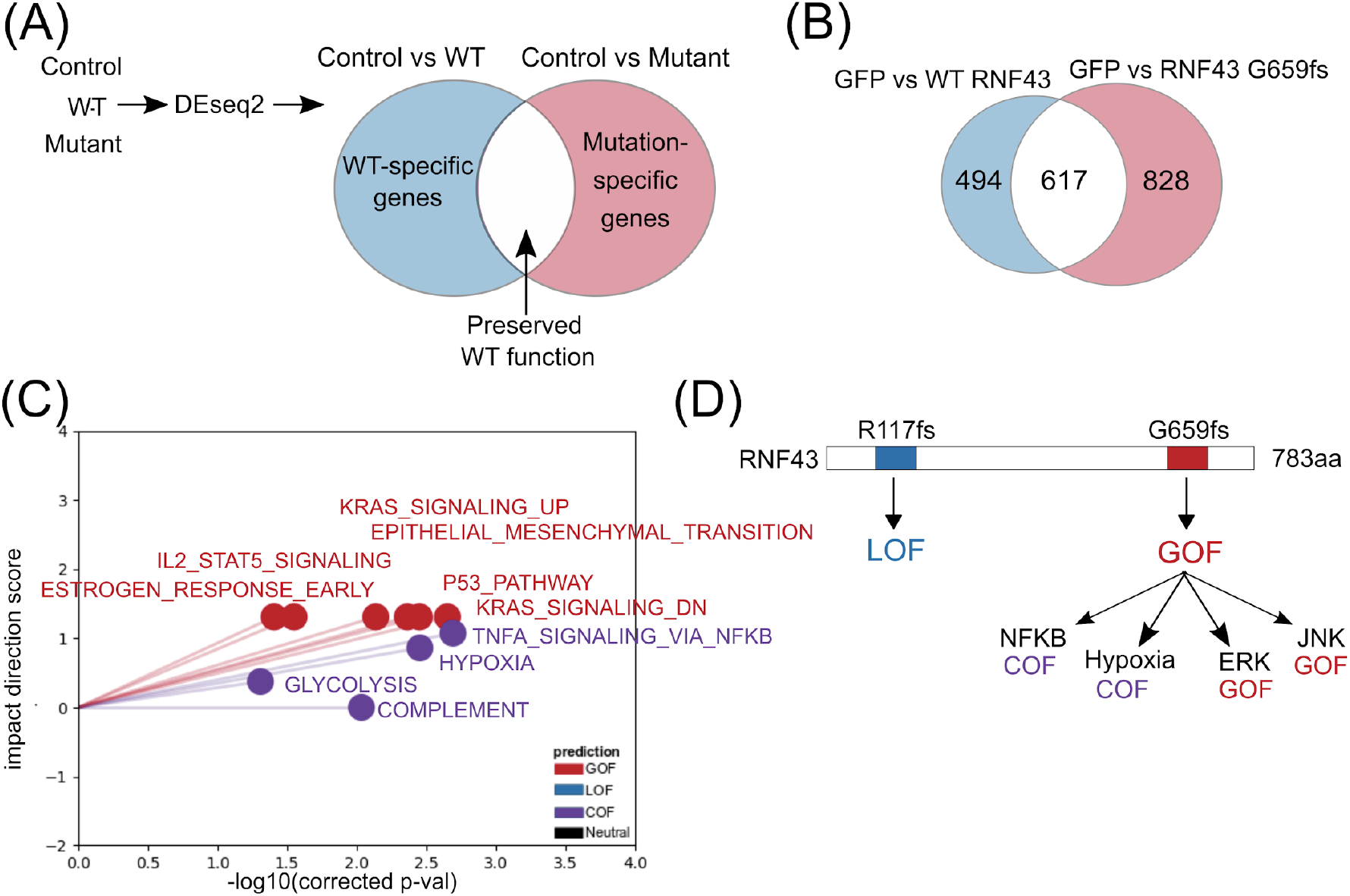
eVIP mutation-specific and WT-specific pathway analysis for GOF *RNF43 G659fs*. (A) Overview of method to identify WT-specific (blue) and mutation-specific (red) differentially expressed genes using DESeq2 (B) Count of GOF *RNF43 G659fs* WT-specific genes(blue) and mutation-specific genes(red) (C) Sparkler plot representation of eVIP Pathway results on GOF *RNF43 G659fs* mutation-specific genes (D) Model summarizing the predicted impacts of the *RNF43* frameshift variants

The LOF *RNF43 R117fs* mutation had 263 mutation-specific and 827 WT-specific differentially expressed genes (S3 Fig). Having more WT-specific genes is consistent with *RNF43 R117fs* being a LOF mutation. The GOF *RNF43 G659fs* mutation had 828 mutation-specific and 494 WT-specific differentially expressed genes, consistent with it being a GOF mutation (Fig 3B).

### eVIP Pathway analysis identifies TNF⍺ via NFKB, KRAS, and hypoxia as the top hallmark pathways impacted by *RNF43 G659fs*

Next, we used the eVIP Pathways approach on the predicted *RNF43 G659fs* GOF mutation to investigate which specific pathways are impacted, thus giving more information on what the function of the mutation is. To find pathways impacted by *RNF43 G659fs*, eVIP Pathways was run separately on the mutation-specific and WT-specific genes (Fig 3C, S4 Fig). We used the 50 hallmark pathway gene sets from MsigDB, to get an eVIP2 functional prediction of GOF, LOF, COF, or neutral for each tested pathway. Due to the smaller number of gene sets, we chose the 50 hallmark pathways over other databases like KEGG or Reactome for simplicity, but these other gene sets can also be used with eVIP Pathways. In order for a pathway to be characterized with eVIP Pathways, a minimum of 10 genes per pathway is required. For *RNF43 G659fs* 10 of the 50 hallmark pathways were tested for the mutation-specific genes, however, *RNF43 R117fs* had no hallmark pathways with at least 10 mutation-specific genes.

When identifying impacted pathways by *RNF43 659fs* from WT-specific genes, the estrogen response late pathway was predicted LOF (S4 Fig, S5 File). Loss of estrogen receptor beta is seen in malignant colon tissue [18]. The mutation-specific genes had 6 GOF pathways and 4 COF pathways (Fig 3C, S6 File). “TNF alpha signaling via NFKappaB” was the most significant pathway and is predicted to be COF. RNF43 has been linked to positive regulation of NF-κB signaling [19]. The other COF pathways are “complement”, “hypoxia”, and “glycolysis”, none of which have been previously associated with RNF43. However, glycolysis has been connected to the Wnt pathway [20], in which RNF43 is known to act, and hypoxic conditions drive glycolysis [21,22].

*RNF43 G659fs* GOF pathways are “KRAS signaling up” and “KRAS signaling down”, “P53 Pathway”, “epithelial mesenchymal transition” (EMT), “IL2 Stat5 signaling”, and “estrogen response early”. *RNF43 G659fs* has not been previously known to activate these pathways; however, WT RNF43 is a known negative regulator of the Wnt/β-catenin pathway which then regulates the RAS-ERK pathway [23–26]

### Reporter assays validate eVIP2 GOF predictions for *RNF43 G659fs*

To validate the eVIP Pathway predictions for the impact of *RNF43 G659fs* on certain pathways, the Cignal 10-Pathway Reporter Array was used to measure the activity of Wnt, Notch, P53/DNA Damage, Cell Cycle/pRB-E2F, NFκB, Myc/Max, Hypoxia, MAPK/ERK, and MAPK/JNK signaling pathways (S1 Table, S2 Table) on HEK-293T cells co-transfected with empty vector, *RNF43 WT*, or *RNF43 G659fs*. NFκB, hypoxia, MAPK/ERK and MAPK/JNK signaling pathways were upregulated in the presence of *RNF43 G659fs* compared with the control vector, but had no significant differences in RNF43 WT transfected cells (Table 1, S5 Fig). RNF43 WT appeared more effective at inhibiting Wnt, consistent with its role as a suppressor of Wnt-signaling [8]. These results are consistent with the finding that the *RNF43 G659fs* is a gain-of-function mutation. Importantly, each of the four impacted pathways (NFκB, Hypoxia, ERK, and JNK) from the functional validation were identified as impactful with eVIP Pathways. Their matching hallmark pathways (“HALLMARK_TNFA_SIGNALING_VIA_NFKB”, “HALLMARK_KRAS_SIGNALING_DN”, and “HALLMARK_HYPOXIA”) were also the top 3 most significant eVIP pathways.

**Table 1.**
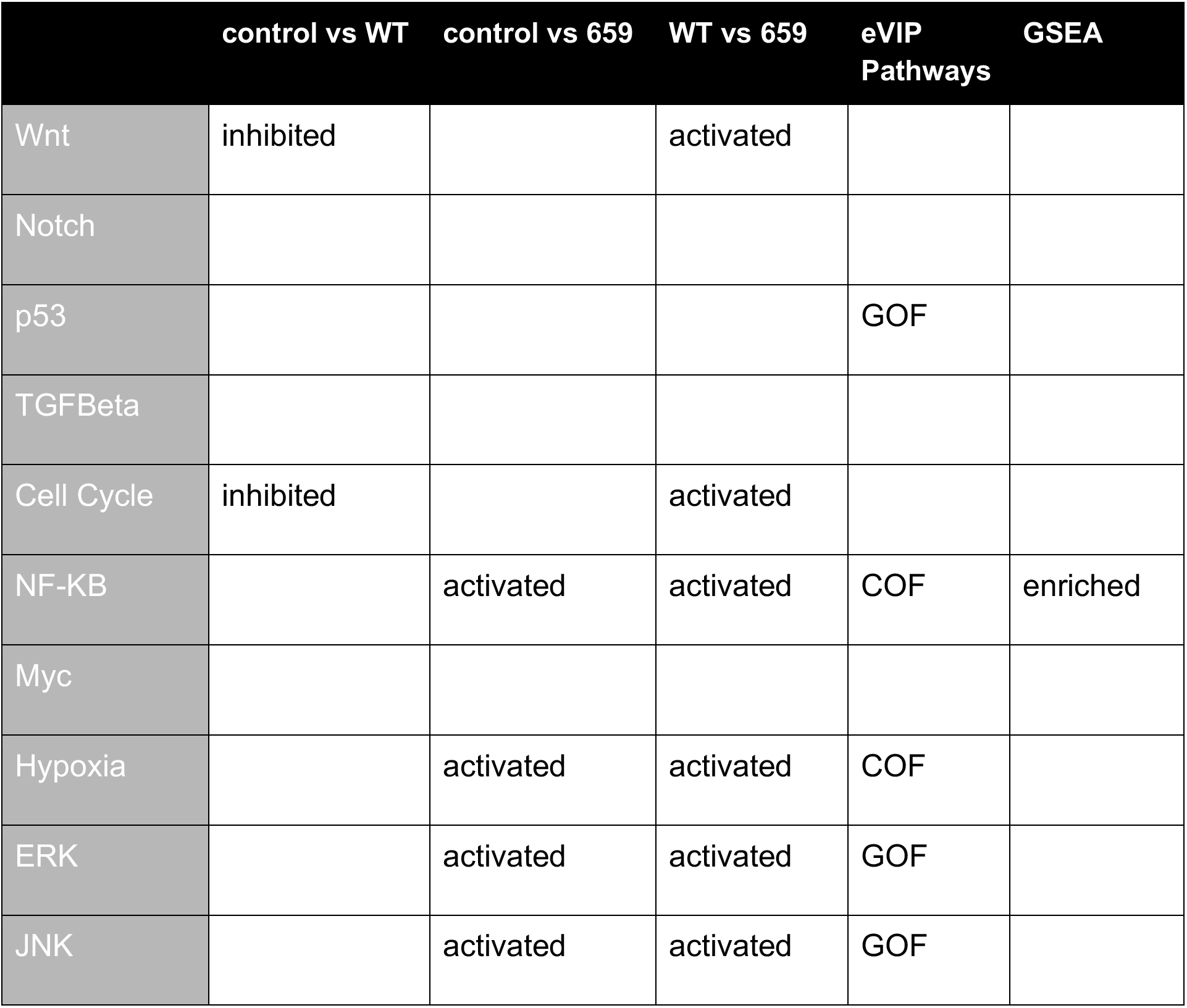
Reporter assay results compared with eVIP Pathways and GSEA Pathway reporter assay results are more consistent with eVIP Pathway prediction than with GSEA. Results from a pathway reporter array using different sample group comparisons (first 3 columns). Pathways were considered significant with p-value under .05, then the direction of difference is reported as inhibited or activated. The last two columns are eVIP Pathway and GSEA results (GOF=Gain of Function, COF = Change of function). For GSEA, a pathway was determined enriched with FDR <.25 and p-value <.05.

|  | control vs WT | control vs 659 | WT vs 659 | eVIP Pathways | GSEA |
| --- | --- | --- | --- | --- | --- |
| Wnt | inhibited |  | activated |  |  |
| Notch |  |  |  |  |  |
| p53 |  |  |  | GOF |  |
| TGFBeta |  |  |  |  |  |
| Cell Cycle | inhibited |  | activated |  |  |
| NF-KB |  | activated | activated | COF | enriched |
| Myc |  |  |  |  |  |
| Hypoxia |  | activated | activated | COF |  |
| ERK |  | activated | activated | GOF |  |
| JNK |  | activated | activated | GOF |  |

The p53 pathway was identified as GOF by eVIP Pathway but was not significant in the assay when compared to the control vector. A possible explanation for this may be in overlap between the mutation-specific genes in the p53 pathway and other pathways. Of the 14 *RNF43 G659fs*-specific differentially expressed genes in the p53 pathway, nine of the genes are in multiple pathways. Specifically, five were also in the “HALLMARK_TNFA_SIGNALING_VIA_NFKB”, “HALLMARK_KRAS_SIGNALING_DN”, and “HALLMARK_HYPOXIA” pathways.

### eVIP Pathways outperforms Gene Set Enrichment Analysis in predicting the three most significantly activated pathways

We compared eVIP Pathways against the popular tool Gene Set Enrichment Analysis (GSEA) [27] using the default parameters on unfiltered gene TPM counts, as recommended. GSEA was run comparing the *RNF43 G659fs* against WT RNF43, resulting in no significant pathways (FDR under 0.25). When comparing *RNF43 G659fs* against GFP, “TNFalpha via NFKB”, apical surface, estrogen response late, EMT, and angiogenesis pathways were significantly enriched (Table 1, S7 File). eVIP and GSEA both identified TNF alpha via NFKB, estrogen late response, and EMT. Though, eVIP Pathways predicts estrogen late as a LOF pathway, adding additional information over GSEA. Notably, GSEA failed to identify the three most activated pathways in the validation assay.

While GSEA and eVIP Pathways both aim to identify biologically meaningful pathways using gene expression changes, they are answering different questions. eVIP Pathways finds pathways that are significantly different in the mutation compared to the WT by incorporating replicate consistency and signature identity. eVIP Pathways looks for consistent changes with large magnitude and high correlation across replicates. GSEA looks for pathways that show cumulative changes in gene expression that are associated with a condition. Whereas eVIP considers each gene set separately, GSEA first ranks all genes by their expression differences and uses that ranking to calculate enrichment scores for each gene set. In contrast to eVIP Pathways, in GSEA, genes that are strongly correlated or anticorrelated with a phenotypic class are given more weight. Our validation results suggest eVIP Pathways is better at predicting which pathways are specifically impacted by a mutation; however, additional studies on other genes and gene variants will need to be performed for a more robust evaluation.

## Discussion

Previous work showed that high-throughput expression-based phenotyping can accurately distinguish between neutral and impactful mutations[5]. eVIP was used to functionally profile a diverse set of 194 lung adenocarcinoma alleles from 53 genes in a single assay, addressing the challenge of interpreting the millions of mutations that have been identified in cancer. An additional strength of the eVIP method is that it can be applied to any mutation, and does not require prior knowledge of the gene. In this study, we present the eVIP2 software, demonstrate that eVIP can be used with RNA-seq data and present eVIP pathway analysis.

Here, we characterize overall and pathway-specific impact of two common frameshift variants in colorectal, gastric and endometrial cancers. Both of the tested frameshift mutations in *RNF43* have been assumed to be loss of function [28]. We show that the two frameshift mutations actually have different effects on RNF43 gene function. While *RNF43 R117fs* was LOF, eVIP2 predicted the *RNF43 G659fs* variant to be a GOF mutant (Figs 2B,C).

We validated *RNF43 G659fs* GOF status through a functional assay, showing the mutant affects the NFκB via TNF alpha, hypoxia, MAPK/ERK and MAPK/JNKpathways, which are not affected by the overexpression of WT RNF43. Importantly, the four pathways differentially affected by *RNF43 G659fs* in the functional assay were identified as COF or GOF with eVIP Pathways and also were the most confident eVIP Pathways predictions. In contrast, GSEA, a common approach to investigate gene sets or pathways affected by a condition, identified only 1 of the 4 significant pathways from the functional assay as enriched.

A recent study claims *RNF43 G659fs* is a passenger mutation, based on its effects on the Wnt pathway [29]. This is consistent with this study, where we did not find the Wnt pathway to be impacted by *RNF43 G659fs* by either eVIP Pathway analysis nor in our validation. However, Tu et. al. mainly focused on the effect on the mutations involvement in the Wnt pathway. With eVIP Pathways, we can profile multiple pathways at once, which is especially helpful when investigating mutations in genes that are not well characterized. We found that *RNF43 G659fs* has a functional impact on other pathways and is unlikely to be a passenger mutation.

The eVIP2 software uses gene expression data from L1000 expression profiling or RNA-seq to predict overall mutation and pathway impact. eVIP Pathways is flexible and can be used with custom gene sets or from existing gene sets from MsigDB, KEGG, or Reactome. Since the original description of the eVIP algorithm, we have improved the software to be more easily run by others to perform similar analyses, thus making this approach more available for mutation profiling by others in the scientific community.

## Availability and future directions

eVIP2 is implemented in Python. The software, instruction manual, tutorial, and example data are available on GitHub (https://github.com/BrooksLabUCSC/eVIP2). For eVIP Pathways, transcript quantification outputs from Kallisto [7] are required as input; however, any molecular profiling data (e.g. L1000, pre-processed gene expression) can be used as input for eVIP overall functional predictions. For future versions of the software, we will test eVIP2 on other molecular profiling such as alternative splicing signatures to investigate the effects of cancer-associated variants.

## Methods

### RNF43 variant functional impact prediction

Quadruplicate transfections of GFP, WT *RNF43*, *RNF43 R117fs*, and *RNF43 G659fs* were done in HEK-293T cells and sequenced with NextSeq 500 (75 nucleotide reads, single end). Supplement Figure 3 showing validation of WT and variant expression was made using Integrative Genome Viewer [30]. Transcript counts were generated using Kallisto [7]. The Kallisto index was built from Ensembl release 94 GRCh38 cDNA transcriptome. Kallisto counts were imported to DESeq2 using tximport and DESeq2 was run using default parameters [11]. Gene set enrichment analysis was performed in python with GSEAPy [27].

### Western blot analysis

HEK293T cells infected with *RNF43* WT, R117fs or G659fs fusion with V5-tag were harvested in RIPA Buffer (Sigma Aldrich, Cat. No. #R0278) supplemented by Protease Inhibitor Cocktail (Cell Signaling, Cat. No. #5871), then resolved by 10% SDS-PAGE. Western blot analysis was performed by the standard method. The protein expression of RNF43 WT and mutants was detected by the primary antibodies anti-V5 Tag (1:5,000, mouse, Monoclonal, Life Technologies, Cat. No. R96025) and anti-β-actin (1:2,000, rabbit, polyclonal, Cell Signaling, Cat. No. #4970) was used as a loading control. Goat anti-Mouse and goat anti-rabbit secondary antibody were obtained from Licor and used at 1:15000 dilution. The proteins of interest were visualized using a two-color Li-COR Odyssey Imager (LI-COR).

### L1000 correlation methods comparison

For the comparison of L1000 weighted connectivity score versus spearman correlation, eVIP was run using default parameters. The plot in Figures S1 uses the corrected p-value (“wt_mut_rep_vs_wt_mut_conn_c_pval”) from the Kruskal-Wallis Test.

### Cignal finder cancer 10-pathway reporter Array

The Cignal pathway reporter assay (SABiosciences/Qiagen, Frederick, MD, USA, Cat. No. CCA-001L/336821) was performed following the instructions provided by the manufacturer. Briefly, 4000 cells/well of HEK-293T cells were seeded in 96-well plates and allowed to settle overnight in a 37°C incubator with 5% CO_2_ before transfection. 100 ng of each Cignal dual-luciferase reporter constructs with 200 ng of RNF43 variants or empty vector constructs were co-transfected into the cells by using Lipofectamine LTX (Thermo Fisher Scientific, Inc.). After 48 hours transfection, cells were harvested and measured the dual-luciferase activities based on Firefly-to-Renilla luminescence ratio using the Dual-Luciferase Reporter Assay System (Promega, Madison, WI, USA).

## Acknowledgements

We would like to acknowledge April Lo and Roman Reggiardo for testing eVIP2. A.M.T was funded through NIH grant 5T32HG008345, the Eugene Cota-Robles Fellowship, and the Bill H. James foundation scholarship. M.G. is funded through a Conquer Cancer Foundation of ASCO Career Development Award, the Cancer Research UK C10674/A27140 Grand Challenge Award and a Stand Up to Cancer Colorectal Cancer Dream Team Translational Research Grant (Grant Number: SU2C-AACR-DT22-17). Stand Up to Cancer is a division of the Entertainment Industry Foundation. Research grants are administered by the American Association for Cancer Research, a scientific partner of SU2C. This work was also partially supported by the Damon Runyon Cancer Research Foundation to A.N.B.

## Author contribution

Conceptualization, A.N.B., A.H.B, M.G., A.M.T.; Data Curation, A.M.T., A.N.B.; Formal Analysis, A.M.T, A.N.B.; Funding Acquisition, A.N.B, A.M.T., M.G.; Investigation, L.F., A.M.T., M.G.; Methodology, A.N.B., A.H.B, A.M.T, M.G.; Software, A.M.T and A.N.B; Supervision, A.N.B.; Validation; L.F.; Visualization, A.M.T; L.F.; Writing – Original Draft Preparation, A.M.T, A.N.B; and Writing – Review & Editing, all authors

---

**S1 Fig.**
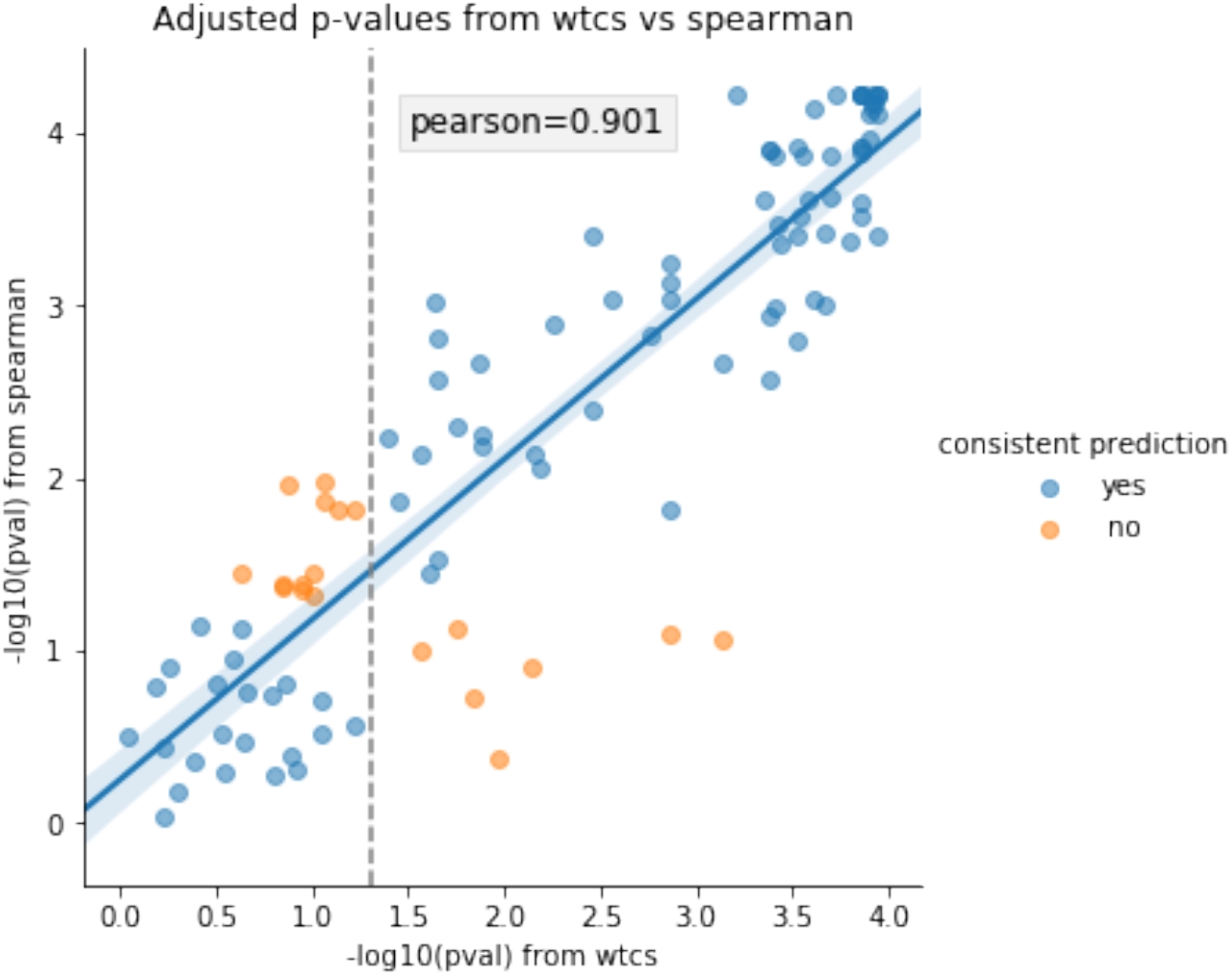
Comparing eVIP correlation metrics. Comparison of eVIP p-values when using spearman rank correlation values or weighted connectivity scores (wtcs) as input. Blue and orange points represent variants that had the same or different impact call with either method, respectively. The dotted vertical line represents p-value cutoff of .05.

**S2 Fig.**
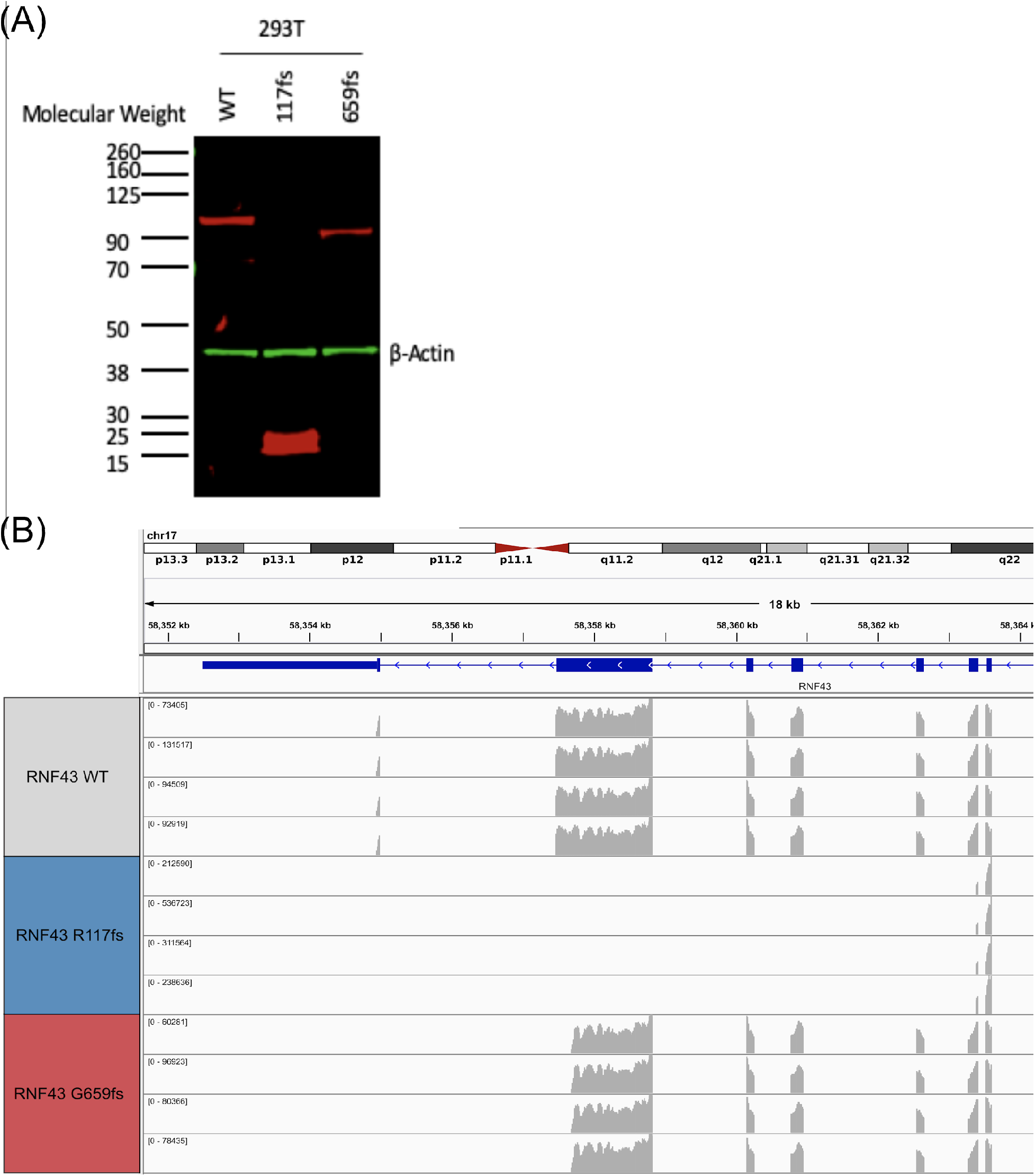
Validation of expression of RNF43 frameshift variants. (A) HEK293T cells infected with RNF43 WT and mutants were verified by western blotting. V5 antibody (Red) indicated RNF43 overexpression, β-Actin (Green) used as control. (B) Expression of RNF43 WT, RNF43 R117fs, and RNF43 G659fs from RNA-Seq in the Integrative Genome Viewer[30]

**S3 Fig.**
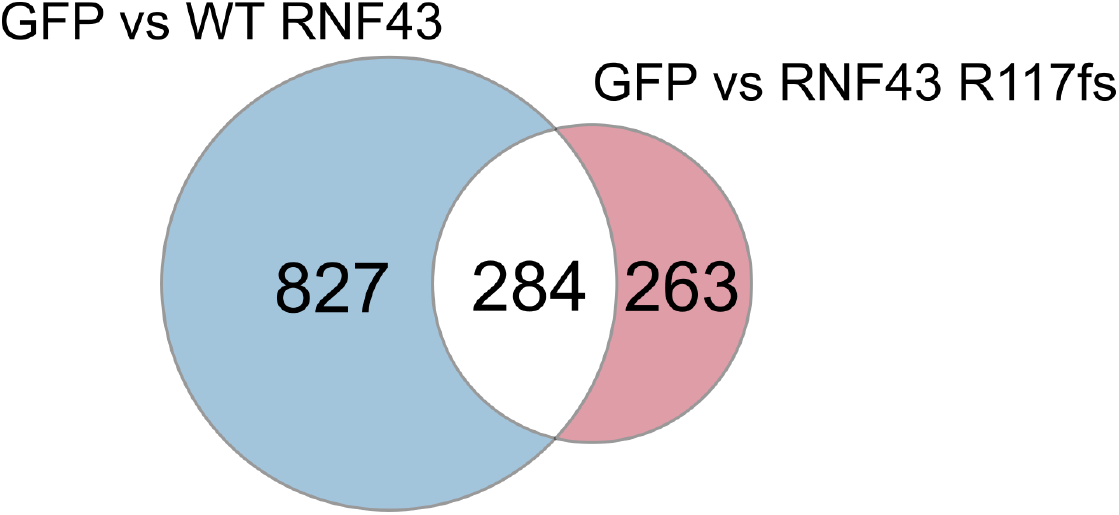
Venn-diagram representation of differentially expressed genes in *RNF43 R117fs*. Count of LOF *RNF43 R117fs* WT-specific genes(blue) and mutation specific genes(red)

**S4 Fig.**
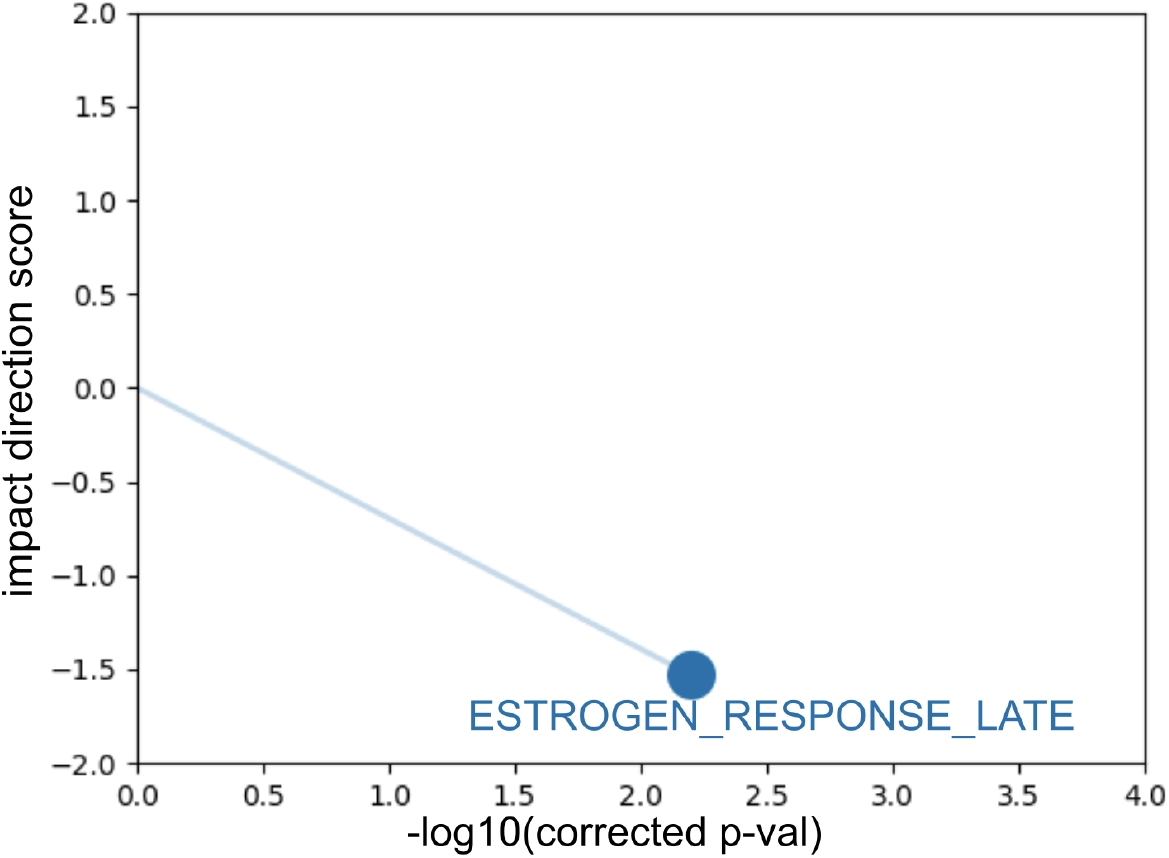
WT-specific Estrogen response late LOF. Sparkler plot representation of eVIP Pathways results on GOF *RNF43 G659fs* WT-specific genes

**S5 Fig.**
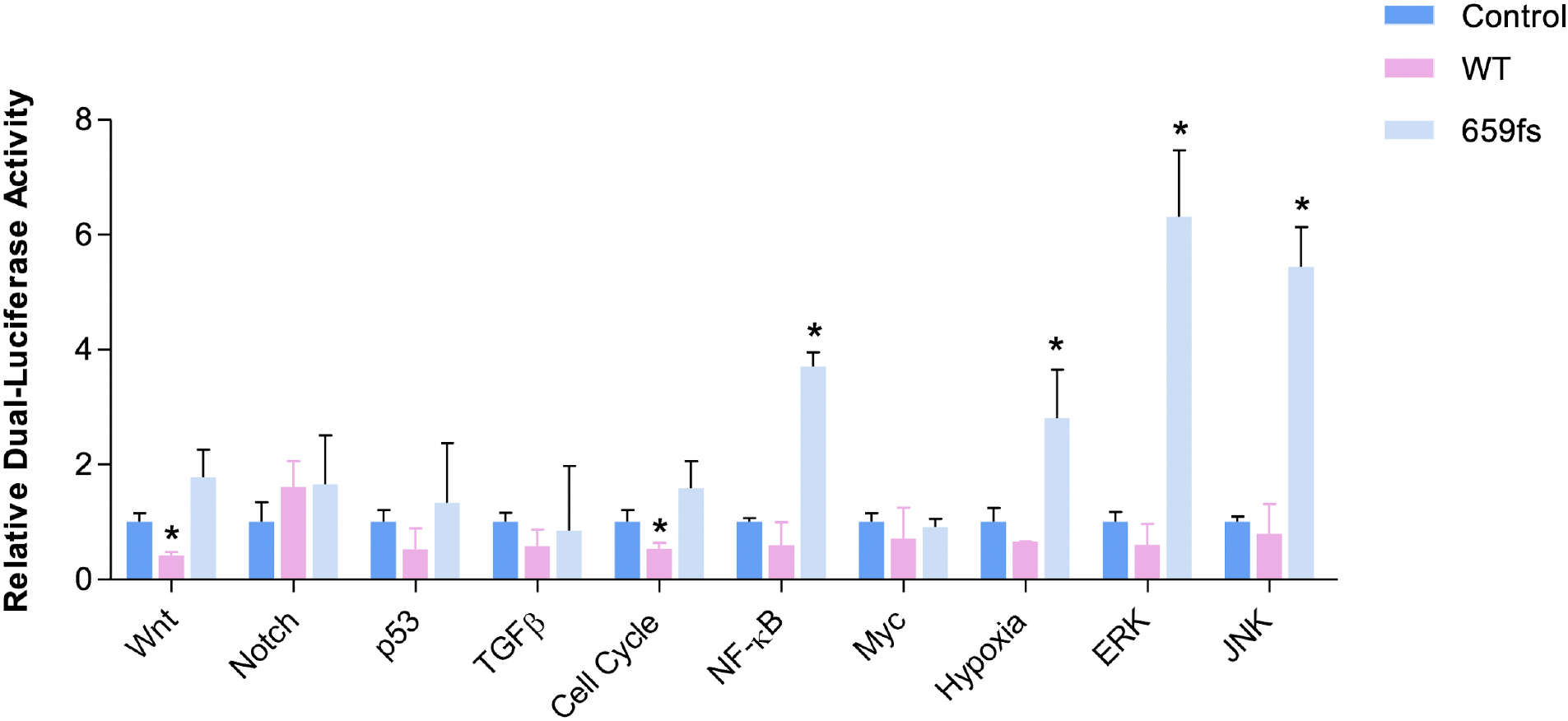
10-Pathway Reporter Array. Cignal 10-Pathway Reporter Array data obtained in HEK-293T cells upon overexpression of Vector, RNF43-WT or RNF43-659fs. Each bar represents the mean±s.d. acquired from three independent experiments. A two-tailed Student’s t-test was used for statistical analysis (*P<0.05).

**S1 Table.**
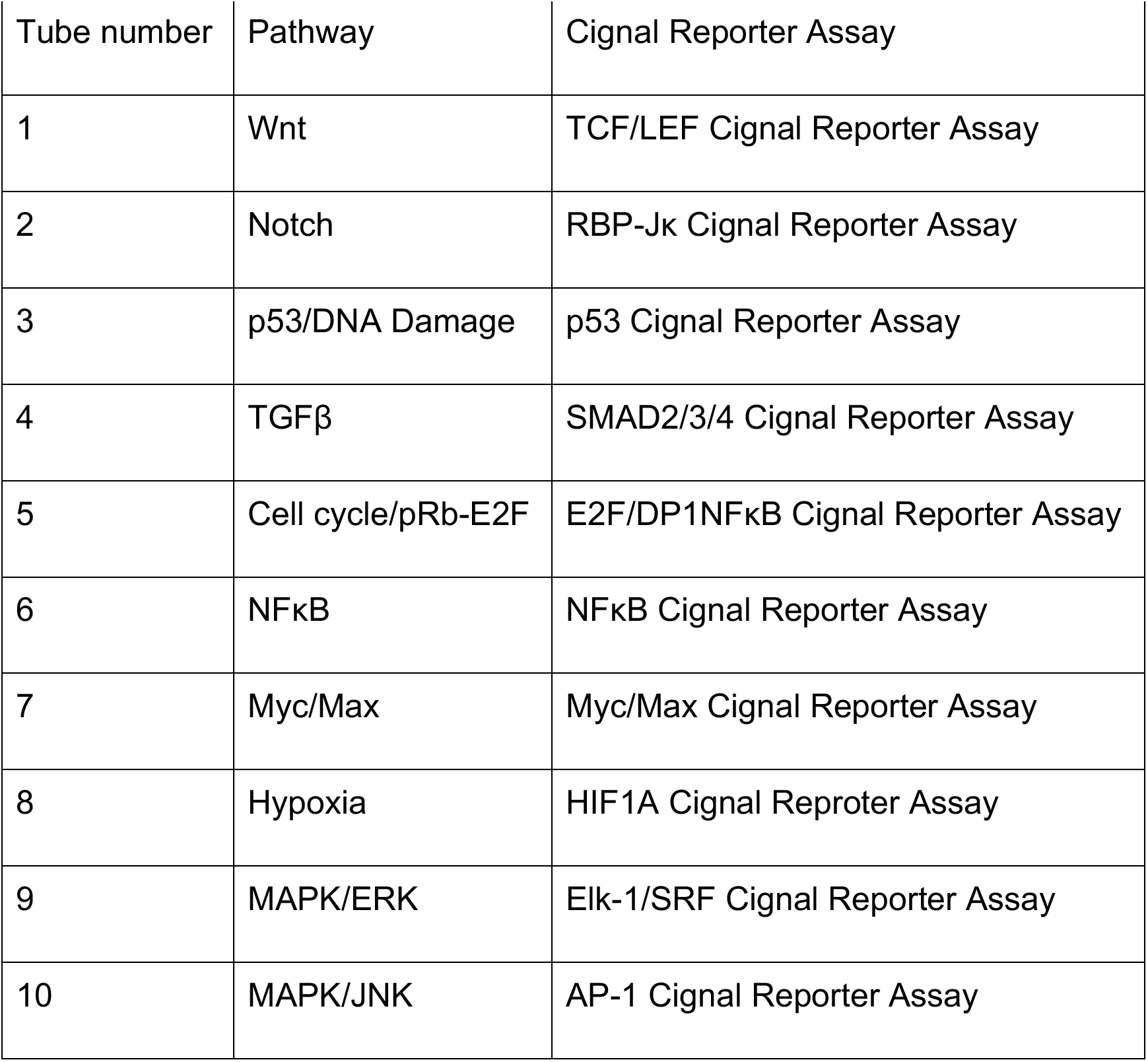
Cignal Reporter Assay used to measure pathway activity

**S2 Table.**
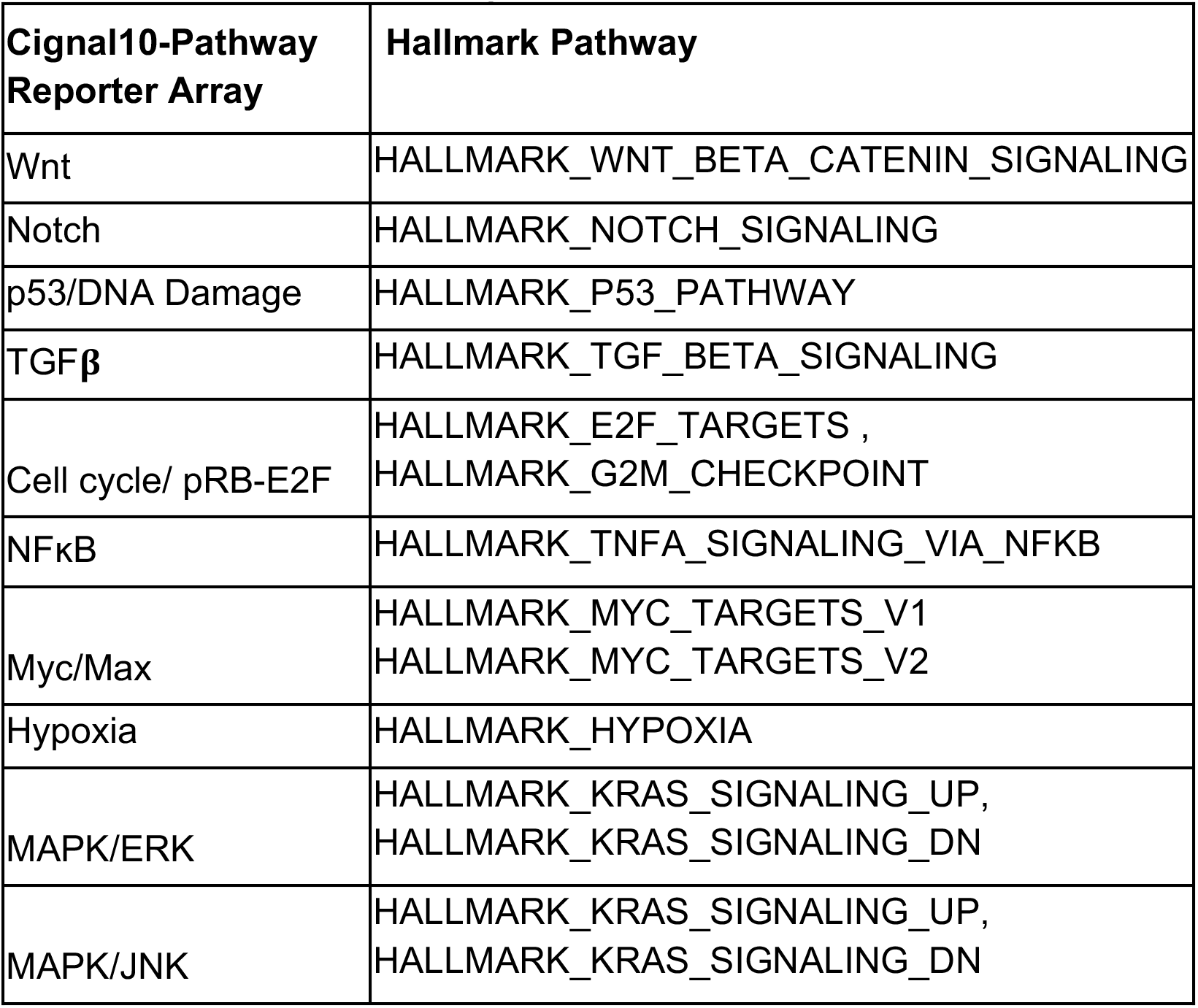
The pathways tested in the Cignal 10-pathway reporter array and their associated hallmark pathways.

## Supplementary Files

**S1 File. Overall eVIP2 outputs**. Text file containing eVIP2 overall functional predictions for *RNF43 R117fs* and *RNF43 G659fs*

**S2 File. DESeq2 GFP vs *RNF43 WT.*** Text file containing DESeq2 outputs for GFP replicates vs *RNF43 WT* replicates

**S3 File. DESeq2 GFP vs *RNF43 R117fs.*** Text file containing DESeq2 outputs for GFP replicates vs *RNF43 R117fs* replicates

**S4 File. DESeq2 GFP vs *RNF43 G659fs.*** Text file containing DESeq2 outputs for GFP replicates vs *RNF43 G659fs* replicates

**S5 File. eVIP Pathways WT-specific genes for *RNF43 G659fs.*** Text file containing eVIP Pathways results for RNF43 G659fs using WT-specific differentially expressed genes

**S6 File. eVIP Pathways mutation-specific genes for *RNF43 G659fs.*** Text file containing eVIP Pathways results for RNF43 G659fs using mutation-specific differentially expressed genes

**S7 File. GSEA results for GFP vs *RNF43 G659fs*.** Text file containing GSEA outputs from GFP replicates vs *RNF43 G659fs* replicates

